# Distinct impacts of sodium channel blockers on the strength-duration properties of human motor cortex neurones

**DOI:** 10.1101/2025.04.16.648964

**Authors:** Lorenzo Rocchi, Kate Brown, Alessandro Di Santo, Hannah Smith, Angel V. Peterchev, John C Rothwell, Ricci Hannah

**Author notes:** Corresponding author: Dr. Ricci Hannah, Centre for Human & Applied Physiological Sciences King’s College London, London, UK.

## Abstract

Voltage-gated sodium channels (VGSCs) are essential for regulating axonal excitability in the human brain. While sodium channel blockers are known to modulate neuronal excitability, their in vivo effects on the human cortex remain poorly understood. Here, we employed novel transcranial magnetic stimulation (TMS) measures to investigate the effects of sodium channel blockers, carbamazepine and lacosamide, on the strength-duration behaviour of human cortical neurones, serving as an index of sodium channel function. Our data showed that single doses of both medications elevated resting motor thresholds compared to placebo, indicating reduced excitability; however, their impacts varied according to TMS pulse width. Carbamazepine raised thresholds proportionally across all pulse widths, whereas lacosamide disproportionately influenced thresholds for long-duration pulses. Crucially, lacosamide reduced the strength-duration time constant and increased rheobase, while carbamazepine had minimal effects on both. These results reveal subtle and unexpected differences in cortical neurone behaviour following VGSC-blocking medication administration. Lacosamide’s response aligns with the proposed mechanism of sodium conductance blockade, while carbamazepine’s effects suggest distinct VGSC interactions or potential off-target effects. Our findings advance the understanding of VGSC-blocking medication interactions in the human cortex and underscore the importance of employing specific TMS measures to gain deeper insights into medication mechanisms of action in vivo. Such measures could serve as valuable adjuncts in medication development and patient monitoring, enhancing understanding of medication action in clinical settings.

## 1. INTRODUCTION

Voltage-gated sodium channels (VGSCs) are critical for neuronal electrical signalling and regulation of axonal excitability, with widespread expression in the brain and peripheral nervous system^1,2^. Rare genetic mutations affecting VGSC function can lead to seizure disorders^3^, underscoring their importance in the cerebral cortex. Accordingly, sodium channels blockers are a first line treatment in some forms of epilepsy^4^. However, despite their clinical effectiveness, and well-documented impact on neuronal excitability and firing characteristics in animal models and *in vitro* [for reviews, see^5,6^], their *in vivo* effects on the human brain remain poorly understood due to the lack of non-invasive tools for assessing ion channel function. This knowledge is vital for understanding therapeutic mechanisms and addressing inter-individual variability in medication efficacy^4^.

Transcranial magnetic stimulation (TMS) is a non-invasive technique for assessing human cortical excitability. A magnetic pulse applied to the primary motor cortex (M1) activates excitatory synaptic inputs to corticospinal neurones, eliciting a muscle twitch on the opposite side of the body^7,8^. This twitch is quantified via electromyography (EMG) as the motor evoked potential (MEP). MEPs are used to evaluate motor threshold—the minimum TMS intensity required to produce an MEP of a specific amplitude—or input–output (I-O) curves, which describe how MEP amplitude changes with varying stimulus intensities^9^. Previous TMS studies have shown that sodium channel blockers like carbamazepine and lacosamide increase motor thresholds and cause a rightward shift in I-O curves^10–12^. These changes were assumed to be caused by the blockade of VGSCs in the neuronal membrane.

However, interpreting the results of medication interventions can be complex. Firstly, different medications with distinct mechanisms of action can lead to the same outcome. Medications that interfere with synaptic transmission at excitatory and inhibitory synapses can also produce changes in motor thresholds^13–16^. Secondly, certain medications may have multiple actions. For instance, carbamazepine may affect gamma-aminobutyric acid (GABA) receptor conductance, calcium channel function, and serotonin release^17–19^. Hence, single motor threshold measurements lack specificity for identifying changes in axonal excitability, often necessitating complementary measures of intracortical excitation and inhibition, presumed to examine synaptic transmission^20^, to rule out ‘off-target’ effects.

Here, we sought to confirm the sodium channel-blocking action using a complementary measure of sodium channel function: the strength–duration time constant (SDTC), which reflects how threshold varies with stimulus strength (amplitude) and duration (width)^21^. The SDTC is thought to reflect the dynamics of VGSCs^22–24^, and prior studies in humans and rodents have shown that sodium channel blockers like lacosamide^25^, mexiletine^26^ and ranolazine^27^ reduce SDTC in peripheral nerves. Our goal was to determine if neurones in the cortex exhibit similar responses, elucidating potential changes in axonal excitability driven by sodium channel blockade.

We used a TMS device capable of varying pulse width^28^, which enables measurement of the SDTC of motor cortical neurones^29–32^, to investigate the effects of single doses of carbamazepine and lacosamide in healthy individuals. These medications were chosen due to their distinct effects on sodium channel function and neuronal firing, as evidenced in animal and *in vitro* experiments^6^.

We tentatively predicted they would produce divergent effects on the strength-duration behaviour. Despite differences in sodium channel sub-type distribution between the cortex and periphery^33^, most sodium channel blockers act non-selectively. We therefore hypothesised that cortical effects would align with those previously observed in peripheral nerves.

## 2. MATERIALS AND METHODS

### 2.1. Participants

Thirteen healthy, young adults (mean age 25 ± 5 years, 7 females) volunteered, providing written informed consent. The sample size is consistent with earlier studies demonstrating effects of lacosamide and carbamazepine on cortical excitability^10–12^. Procedures adhered to the Declaration of Helsinki and were approved by the University College London Research Ethics Committee (Project ID: 5732/003). All participants were free from neurological or psychiatric disorders and not taking neuroactive medications.

### 2.2. Experimental design

This placebo-controlled, single-blind, repeated-measures study involved three laboratory visits, during which participants—blinded to treatment condition—received carbamazepine, lacosamide, or placebo in a randomised order. Washout periods lasted at least seven days. TMS measurements were conducted at a single time point corresponding to the presumed peak plasma concentration of the medications: 6 h post-carbamazepine^34,35^ and 1 h post-lacosamide^36^ or placebo. Measurements included a preliminary estimate of resting motor threshold (RMT) and I-O curves.

### 2.3. Medications

The placebo was a lime and mint-flavoured drink, also used to mask the taste of the anti-seizure medications. Carbamazepine (600 mg) and lacosamide (200 mg) were administered in liquid form, with doses selected based on prior evidence of their effects on motor thresholds and I-O curves^10–12^, and peripheral nerve SDTC^25^.

### 2.4. Electromyography (EMG)

EMG was recorded using a bipolar muscle-tendon montage with one electrode positioned over the belly of the right first dorsal interosseous (FDI) muscle and the reference over the second metacarpophalangeal joint, with the ground electrode placed on the ulnar styloid process. EMG signals were amplified (×1000; Digitimer D360, Digitimer, UK) and bandpass filtered (5–3000 Hz). They were digitised at 5000 Hz (Power1401, Cambridge Electronic Design, UK) and analysed with Signal software version 5.10 (Cambridge Electronic Design, UK).

### 2.5. Transcranial magnetic stimulation (TMS)

TMS was delivered to the left M1 via a device connected to a 70 mm figure-of-8 coil (Elevate TMS, Rogue Research Inc., Canada) to evoke MEPs in the right FDI muscle. The induced electric field waveform (Figure 1) had a pseudo-rectangular shape with an M-ratio of 0.2 (ratio of the amplitude of the initial phase to that of the second phase), favouring unidirectional electric field pulses. We used three different pulse widths (30, 60 and 120 μs duration of the initial phase), with the maximum stimulator output (MSO) limited to 100%, 73% and 50%, respectively.

**Figure 1.**
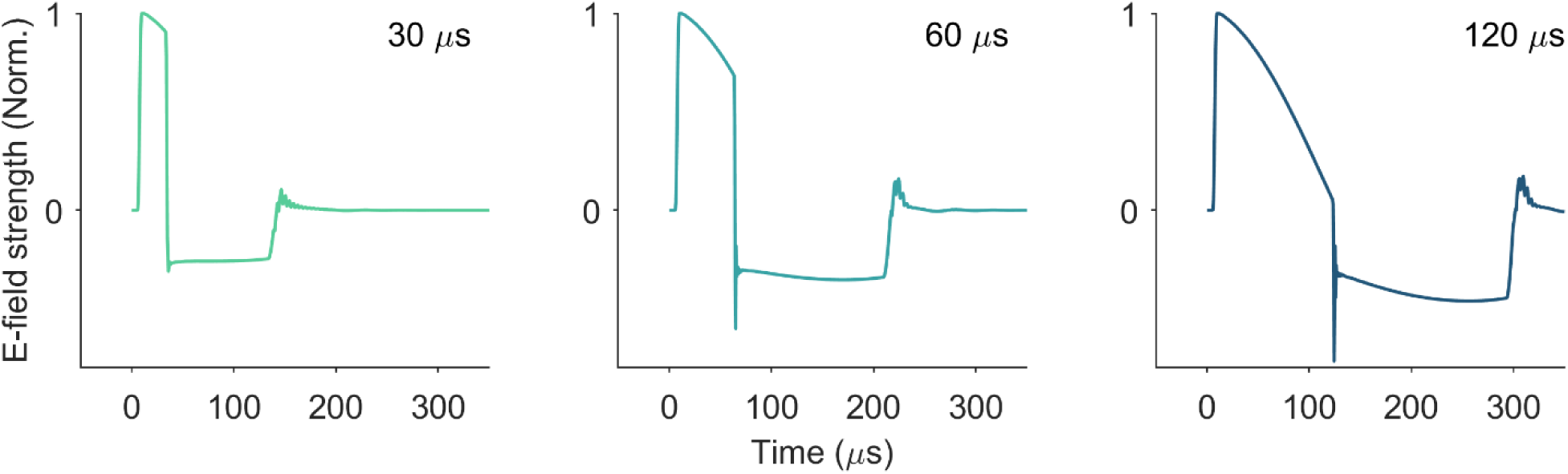
Electric field waveforms recorded with a search coil for each of the three pulse widths of the initial phase (30, 60 and 120 μs). Amplitudes are normalised to unity.

### 2.6. Experimental protocol

Participants received the medication or placebo and waited/returned to the lab 1 h (placebo, lacosamide) or 6 h (carbamazepine) before TMS procedures. Participants were then seated. The TMS coil was positioned over the left M1 with the handle angled ∼45° postero-laterally to induce posterior–anterior initial currents with respect to M1. The motor hotspot was identified by locating the position where slightly suprathreshold 120 μs pulses elicited the largest and most consistent FDI MEPs, and marked on a participant’s cap.

Preliminary RMT estimates for each pulse width were determined at the hotspot by finding the intensity that evoked MEPs > 0.05 mV in 5 of 10 trials. These values guided stimulus intensities for I-O curves, which provided more robust RMT estimates and a comprehensive description of cortical excitability in response to the medications. Ten pulses were delivered at 11 intensities for each pulse width: 80, 90, 100, 108, 116, 124, 132, 140, 148, 156 and 164% preliminary RMT. Within each pulse width, the order of stimulus intensities was also randomised.

### 2.7. Data analysis

The strength–duration behaviour of nodal membranes can be described by two parameters: the SDTC, describing the relationship between the strength (amplitude) and duration (width) of a pulse needed to elicit an action potential, and the rheobase, the extrapolated threshold for a pulse of infinite duration^22,37^. As in previous TMS studies, we employed the RMT to characterise the strength-duration behaviour^30,32^.

#### 2.7.1. Estimating resting motor thresholds from input-output curves

We used the I-O curve to derive a more precise RMT estimate (RMT_I-O_). For each pulse width, ten stimuli were delivered per intensity, with values log-transformed (log₁₀) and the median response at each intensity calculated. These medians were used to fit a sigmoidal curve (see Figure 2):

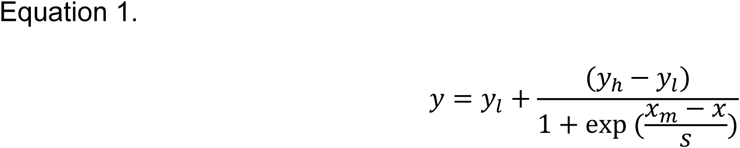

**Figure 2.**
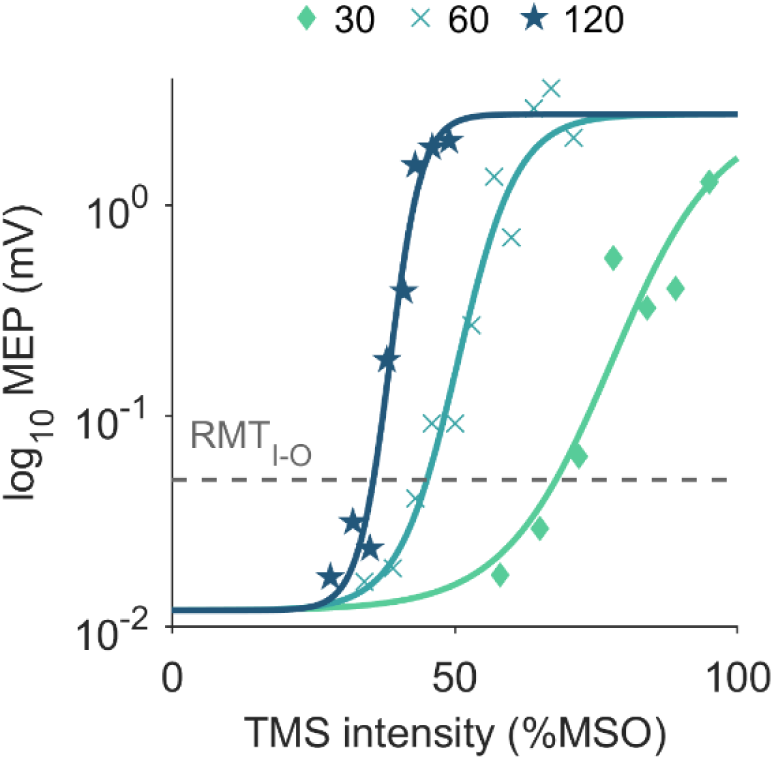
Example input–output (I-O) curves for each pulse width from a single participant. Median log-transformed MEP peak-to-peak amplitude as a function of TMS stimulus intensity. The curves were used to estimate stimulus-response characteristics (*s,* reflecting spread or width of the curve and *x*_*m*_, reflecting the TMS intensity at the midpoint in the curve), as well as RMT_I-O_. The group average model fits, based on the Boltzmann sigmoidal curve, exhibited *R*^2^ values ranging 0.94 to 0.97 across all pulse widths and conditions. Individual I-O curves for each session are provided in Supplementary Figures 1-3.

Here *x* is the stimulus intensity (%MSO), *y* is the log-transformed MEP amplitude (mV), *y*_*l*_ and *y*_ℎ_ are the lower and upper saturation levels, *x*_*m*_ is the stimulus intensity at the midpoint of the curve (%MSO), and *s* is the spread or width of the curve specifically describing the speed of transition between the lower and upper asymptotes. Note that the inverse of the *s* is directly proportional to the steepness of the slope of the I-O curve at *x*_*m*_.

Saturation levels (*y*_*l*_ and *y*_ℎ_) were fixed across pulse widths within each session. *y*_*l*_ was determined as the 10^th^ percentile of the peak-to-peak EMG amplitudes measured in a 35 ms window prior to TMS, reflecting the background noise. *y*_ℎ_ was set as the 90th percentile of the peak-to-peak MEP amplitudes, measured in a 35 ms window after the TMS pulse (15-50 ms). Both were calculated across all pulse widths and stimulus intensities within a session. Curve fitting adjusted *s* and *x*_*m*_ to optimise the fit of the curve to the data. The parameters from the model were used to estimate RMT_I-O_ as the stimulus intensity producing an MEP amplitude of 0.05 mV after log transformation, i.e. *log*_10_(0.05). We fitted these curves for each pulse width of each experimental condition and participant.

We also determined the slope of the I-O curve (*k*) using linear regression, rather than the *s* parameter from the Boltzmann fit, which could be influenced by occasional incomplete plateauing of the curves (note that this would not affect the estimation of RMT_I-O_ or strength-duration behaviour). As described earlier, saturation levels were fixed across pulse widths, helping to address this issue; however, the approach to calculating *k* sidesteps any uncertainty introduced by non-plateauing curves. We fitted a straight line to the median values of the log-transformed MEP amplitudes and corresponding stimulus intensities. For each pulse width during a session, we applied linear regression across four consecutive intensities, starting at 80% RMT. This process was repeated for subsequent intensity steps, iterating the procedure until the initial intensity reached 140% RMT. We then took the greatest *k* value from these iterations as our final slope estimate. The rationale for this iterative approach was that the linear portion of the curve might vary due to minor imprecisions in the initial RMT estimates and differences in the shape of the I-O curves both within and across individuals.

#### 2.7.2. Strength–duration time constant

We employed similar procedures to those in our previous studies^31,32^, deriving strength–duration curves for each experimental condition using RMT_I-O_ across pulse widths. The SDTC models how stimulus intensity changes with pulse width, which depends on the local electric field and axonal membrane properties. This relationship is described by the equation:

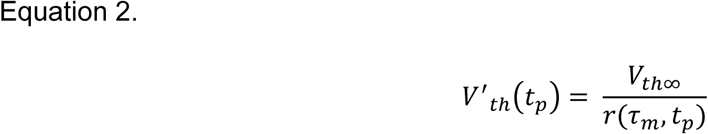

where *V*^′^_*t*ℎ_*(t*_*p*_) is the modelled RMT (the threshold required to evoke a motor response), *V*_th∞_ is the rheobase, *t*_*p*_ is the pulse width, τ_*m*_ is the SDTC, and *r(*τ_*m*_ *t*_*p*_) is a depolarisation factor represents how effectively the TMS pulse depolarises the membrane^32^.

To estimate the rheobase and SDTC, we fitted the parametric model *V*^′^_*t*ℎ_*(t*_*p*_) to the experimental RMT_I-O_ data, *V*_*t*ℎ_*(t*_*p*_). We did this by minimising the difference between the predicted and actual RMT_I-O_ values using the following least-squares method:

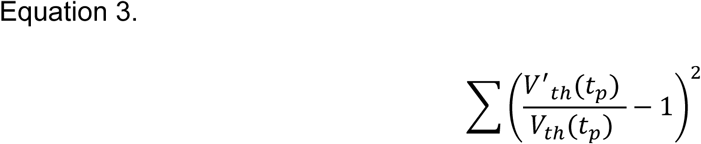

This method helps to find the best-fitting values for rheobase and SDTC by reducing discrepancies between the model and the experimental data. The SDTC and rheobase calculations were performed in Matlab (R2024b, The Mathworks Inc., USA). Parameters were estimated for each participant and each experimental condition under the assumption of individual SDTC and rheobase values.

### 2.8. Statistical analyses

Data are presented as mean and standard error of the mean (SEM). To examine the effects of medication condition and pulse width on RMT_I-O_ and *k* we used linear mixed-effects (LME) models, with ANOVA applied to the fixed effects. This approach accounts for inter-individual variability by including subject-specific random intercepts and slopes, thereby modelling the correlation structure of repeated measures within subjects. The fixed effects tested included medication condition (placebo, carbamazepine, lacosamide), pulse width (30, 60, and 120 μs), and their interaction.

To examine the effects of medication condition on SDTC and rheobase, we used a similar LME approach, including random intercepts to account for subject-specific variability. The placebo condition was treated as the reference category, and the effects of carbamazepine and lacosamide were estimated relative to this reference. SDTC and rheobase are linearly and inversely related^32,38,39^, likely due to their shared dependence on sodium conductance. We therefore expected the medications to exert opposing effects on these indices. Linear regression evaluated the relationship between SDTC and rheobase for each condition. Additionally, an LME model examined this relationship across conditions, including medication condition and their interaction as fixed effects, with random intercepts to account for within-participant variability.

All statistical analyses were conducted with a significance level set at p < 0.05. Effect sizes for fixed effects are reported using Cohen’s *d*, calculated as the fixed effects estimate divided by the residual standard deviation.

## 3. RESULTS

### 3.1. Side-effects

The medications were generally well-tolerated with only mild side-effects including drowsiness (carbamazepine, 9 participants; lacosamide, 7 participants), erythema (carbamazepine, 1), dizziness (carbamazepine, 1), nausea (carbamazepine, 1) and mild ataxia (carbamazepine, 1). None of the side effects interfered with participants’ ability to complete the experiments.

### 3.2. Single motor threshold analysis

We first analysed the impact of each medication on a single threshold, RMT_I-O_ measured at 60 µs (Figure 3A), which is close to a standard ∼80 microsecond pulse used in previous studies. This analysis was performed to establish whether the medications differed from placebo in their effects on motor threshold, as a first step in understanding their impact on cortical excitability. The analysis showed significant main effects for medication (F_[df]_ = 6.20_[2, 36]_, p = 0.005). Fixed effect estimates showed that both carbamazepine (estimate = 6.69, SE = 0.81, t_[df]_ = 3.33_[36]_, p = 0.002, Cohen’s *d* = 1.31) and lacosamide (estimate = 2.15, SE = 0.81, t_[df]_ = 2.66_[36]_, p = 0.012, Cohen’s *d* = 1.04) significantly increased RMT_I-O_ compared to placebo. To directly compare the effects of carbamazepine and lacosamide, we ran a similar LME model with carbamazepine as the reference category. This model showed no significant difference between the effects of carbamazepine and lacosamide (estimate = –0.54, SE = 0.81, t_[df]_ = –0.67_[36]_, p = 0.51, Cohen’s *d* = –0.21). Hence, analysis of single motor thresholds does not distinguish between the effects of the two medications.

**Figure 3.**
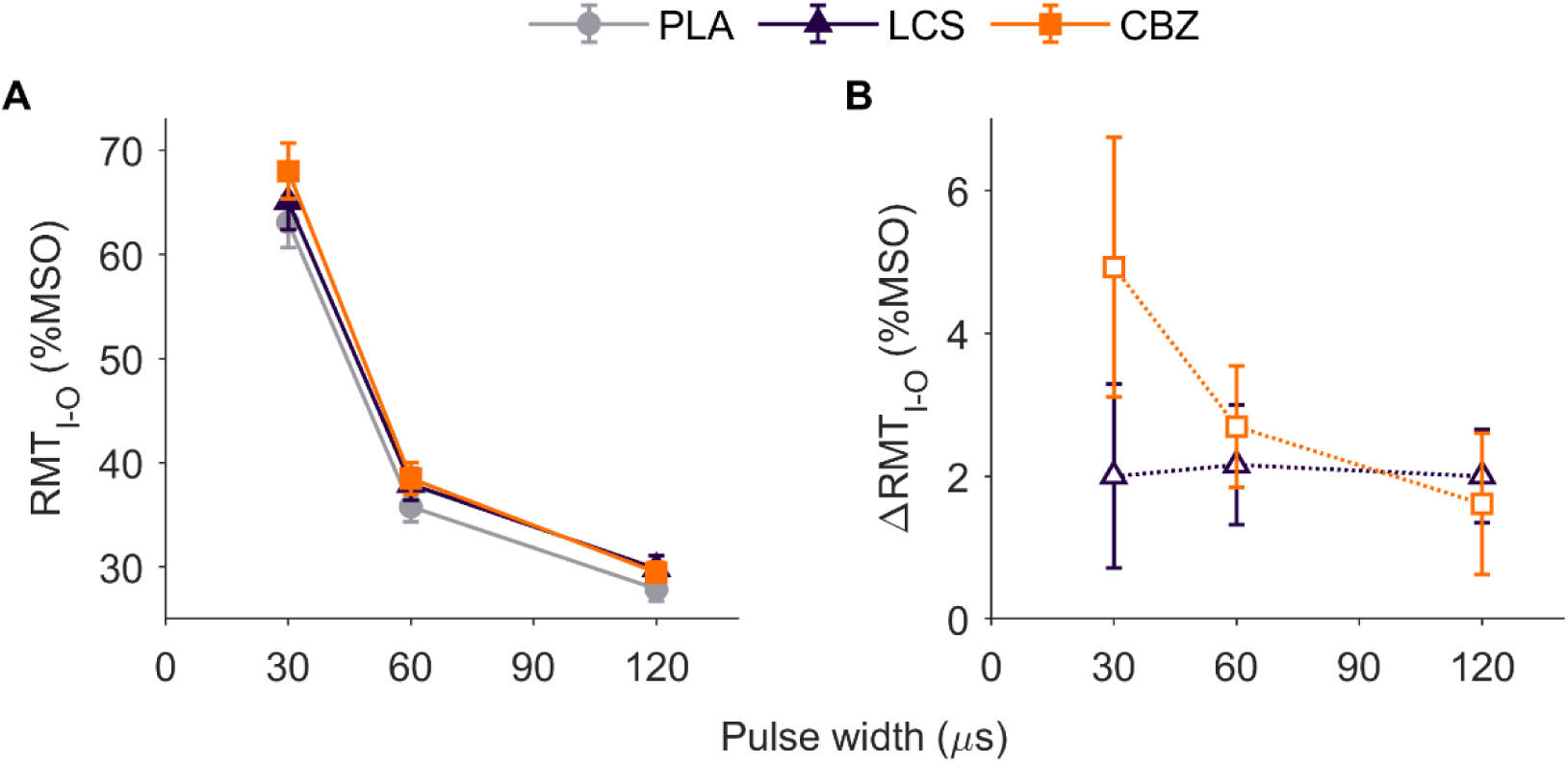
Strength-duration behaviour derived from estimates of RMT_I-O_. **A.** Strength–duration curves of RMT_I-O_ as a function of pulse width (30, 60, 120 μs) for each experimental condition (placebo, carbamazepine, lacosamide). Motor thresholds decrease with increasing pulse width, with carbamazepine and lacosamide conditions showing elevated thresholds compared to placebo. **B.** Differences in RMT_I-O_ between each medication condition (carbamazepine, lacosamide) and placebo across pulse widths. Positive values indicate an increase relative to placebo. The differences for carbamazepine are pulse-width-dependent, scaling with RMT_I-O_ in panel A, whereas differences for lacosamide are consistent across pulse widths. Data are mean ± SEM.

### 3.3. Strength–duration curves

We next explored the strength–duration behaviour by using the I-O curves to estimate RMT_I-O_. Motor thresholds in each experimental condition displayed the expected strength-duration behaviour^31,32^, where RMT_I-O_ was greater for brief versus long duration pulses (Figure 3A). This was evident in the main effect of pulse width in the ANOVA (F_[df]_ = 228.75_[2, 108]_, p = 1.49 ×10^-39^), with the fixed effects showing that RMT_I-O_ with 60 µs (estimate = –27.31, SE = 1.40, t_[df]_ = – 19.47_[108]_, p = 4.17×10^−37^, Cohen’s *d* = –10.9) and 120 µs pulses (estimate = –35.23, SE = 1.70, t_[df]_ = –20.75_[108]_, p = 1.81×10^−39^, Cohen’s *d* = –14.0) were reduced compared to with 30 µs pulses.

Moreover, the results showed that motor thresholds were generally increased in both medication conditions compared to the placebo (Figure 3A). The pattern of results was reflected in the main effect of medication in the ANOVA (F_[df]_ = 12.58_[2, 108]_, p = 1.23×10^−5^). Fixed effect showed that both carbamazepine (estimate = 4.92, SE = 0.99, t_[df]_ = 4.99_[108]_, p = 2.36×10^−6^) and lacosamide (estimate = 2.00, SE = 0.99, t_[df]_ = 2.03_[108]_, p = 0.045) produced significant increases in RMT_I-O_ compared to placebo. The interaction between pulse width and medication was not significant (F_[df]_ = 1.97_[4, 108]_, p = 0.104).

To further examine medication-specific effects, we conducted follow-up analyses using LME models restricted to pairwise comparisons (i.e., placebo versus carbamazepine, placebo versus lacosamide, and carbamazepine versus lacosamide). These analyses included only two medication conditions at a time, enabling direct tests main effects and condition-by-pulse width interactions.

For lacosamide compared to placebo, the LME revealed a significant main effect of condition but no interaction (condition, F_[1,72]_ = 5.58, p = 0.021; pulse width, F_[2,72]_ = 256.6, p = 1.74×10^−33^; interaction F_[2,72]_ = 0.011, p = 0.99), suggesting a consistent absolute increase in RMT_I-O_ across pulse widths. In contrast, carbamazepine versus placebo showed a strong main effect of condition with a trend towards an interaction (condition, F_[1,72]_ = 21.16, p = 1.77×10^−5^; pulse width, F_[2,72]_ = 223.2, p = 1.36×10^−31^; interaction F_[2,72]_ = 2.49, p = 0.091), hinting at pulse-width dependent effects. Finally, a direct comparison between carbamazepine and lacosamide revealed a significant condition-by-pulse width interaction (condition, F_[1,72]_ = 11.88, p = 9.5×10^−4^; pulse width, F_[2,72]_ = 257.7, p = 1.52×10^−33^; interaction F_[2,72]_ = 4.05, p = 0.021). This implies that the effects of carbamazepine were pulse width-dependent, whereas lacosamide elicited more uniform responses across pulse widths.

To visualise this more effectively, we calculated the difference in RMT_I-O_ between each medication conditions and the placebo (Figure 3B). Carbamazepine exhibited pulse-width-dependent variations in absolute RMT_I-O_ differences, with larger shifts at shorter pulse widths where thresholds are higher. Consequently, absolute differences at each pulse width were broadly proportional to the magnitude of RMT_I-O_. In contrast, lacosamide showed consistent absolute differences that were therefore not proportional to RMT_I-O_ across widths. This was supported by the ANOVA on the differences within the same LME framework, which showed an interaction (condition, F_[1,72]_ = 9.82, p = 0.003; pulse width, F_[2,72]_ = 4.24, p = 0.018; interaction, F_[2,72]_ = 3.35, p = 0.041).

### 3.4. Strength–duration time constant and rheobase estimates

The strength–duration curves were then formally described by estimating the SDTC and rheobase. ANOVA on the fixed effects revealed a main effect of condition on the SDTC (F_[df]_ = 3.73_[2, 36]_, p = 0.034, Figure 4A). Fixed effects showed that the SDTC was reduced for the lacosamide condition compared to placebo (estimate = –67.6, SE = 30.4, t_[df]_ = –2.22_[36]_, p = 0.032, Cohen’s d = –0.87). By contrast, there was no statistical difference between the placebo and carbamazepine conditions (estimate = 7.85, SE = 30.4, t_[df]_ = 0.26 _[36]_, p = 0.798, Cohen’s d = – 0.10). A LME model with carbamazepine as the reference category showed that the SDTC was reduced in the lacosamide condition compared to the carbamazepine (estimate = –75.5, SE = 30.4, t_[df]_ = –2.48_[36]_, p = 0.018, Cohen’s *d* = –0.97).

**Figure 4.**
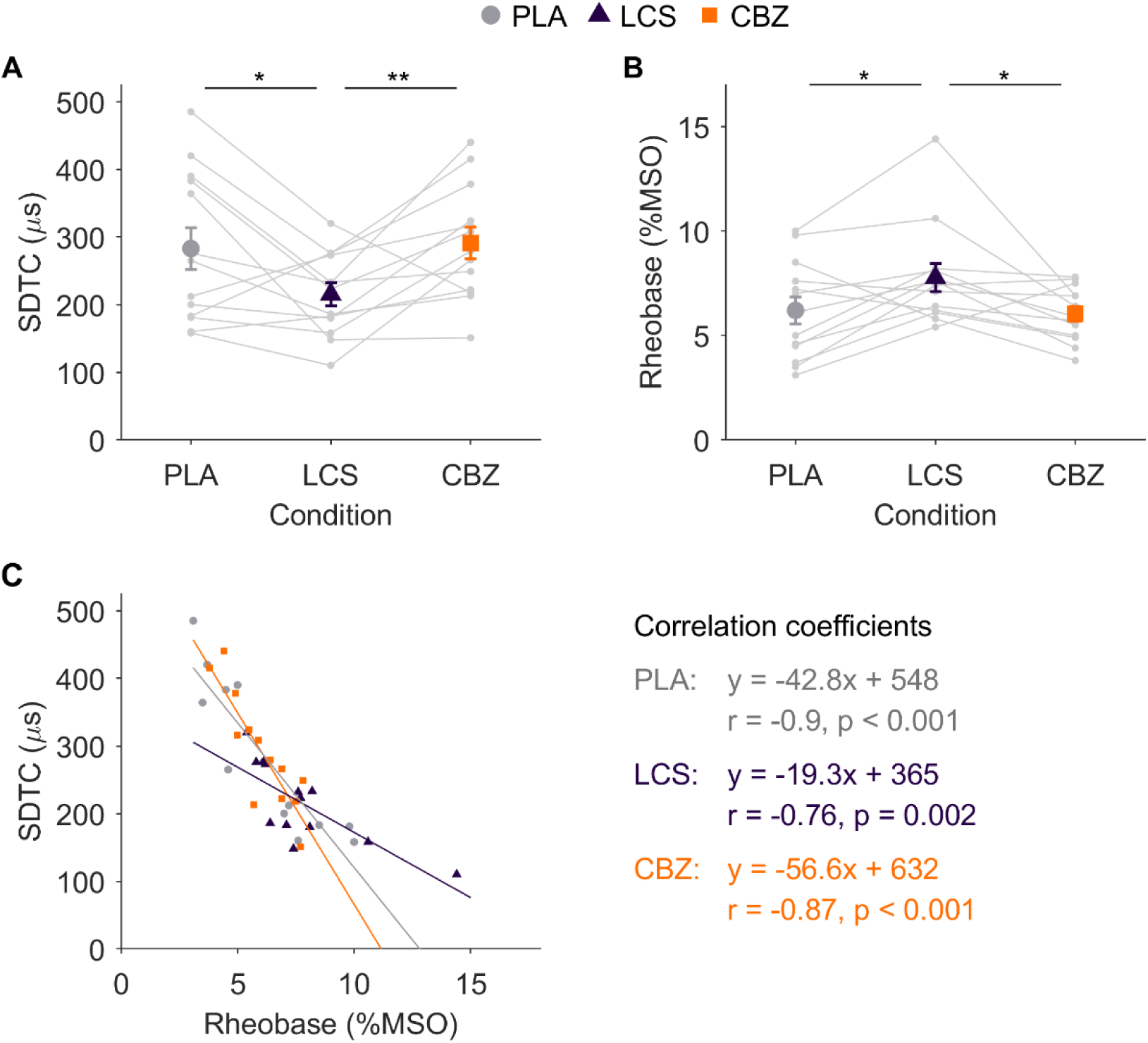
Strength-duration parameter estimates derived from RMT_I-O_ across medication conditions (placebo, carbamazepine, lacosamide). **A.** Strength–duration time constant (SDTC) for each condition. Lacosamide significantly reduced SDTC compared to placebo and carbamazepine, while no difference was observed between placebo and carbamazepine. **B.** Rheobase estimates for each condition. Lacosamide significantly increased rheobase compared to placebo and carbamazepine, while no difference was observed between placebo and carbamazepine. Data in panels A and B are shown as mean ± SEM, with smaller dots representing individual participant data. **C.** Inverse relationship between individual SDTC and rheobase estimates across conditions, with simple linear regression fits shown for each condition. Regression slopes and intercepts hint at differences between conditions, particularly for lacosamide, which demonstrates a less pronounced negative relationship compared to placebo and carbamazepine. Each point represents data from a single participant, with strong correlations observed within all conditions.

For the rheobase, ANOVA on the fixed effects revealed a main effect of condition (F_[df]_ = 4.24_[2, 36]_, p =0.022, Figure 4B). Fixed effects showed that the rheobase was increased for the lacosamide condition compared to placebo (estimate = 1.57, SE = 0.66, t_[df]_ = 2.38_[36]_, p = 0.023, Cohen’s d = 0.93). By contrast, there was no statistical difference between the placebo and carbamazepine conditions (estimate = –0.17, SE = 0.66, t_[df]_ = –0.26 _[36]_, p = 0.799, Cohen’s d = – 0.10). A LME model with carbamazepine as the reference category revealed that the rheobase was greater in the lacosamide condition compared to the carbamazepine (estimate = 1.57, SE = 0.66, t_[df]_ = –2.38_[36]_, p = 0.023, Cohen’s *d* = 0.94).

Further analysis, modelling the data under the assumption of a common SDTC for all participants within the same medication intervention condition (but with individual rheobase values), was conducted to mitigate potential overfitting associated with estimating SDTC separately for each participant^32^. This approach confirmed a similar pattern of results (Supplementary Table S1).

Taken together, the results suggest that lacosamide may have an impact on the strength-duration properties in comparison to both placebo and carbamazepine, manifesting opposing effects on the rheobase and SDTC.

To further explore the seemingly opposing effects on SDTC and rheobase, we evaluated the relationship between the variables for each medication condition. We found that individual estimates of SDTC and rheobase were strongly and inversely related to one another within each condition (Figure 4C; all p < 0.01).

Moreover, the data suggests that the slope of the relationship differs numerically across the three conditions, and this was supported by the results of the LME model. The main effect of rheobase indicates a strong negative relationship (estimate = 46, SE = 3.2, t_[df]_ = –14.0_[33]_, p = 1.8 × 10^-15^), where each 1% increase in rheobase corresponds to an approximate 46 µs reduction in SDTC. The intercept in the model represents the estimated SDTC for the placebo condition (estimate = 566, SE = 23, t_[df]_ = 24.2_[33]_, p = 1.5 × 10^-22^), with all predictors set to their reference levels, while the estimates for lacosamide and carbamazepine reflects their effect relative to this baseline. The model shows a significant main effect of lacosamide (estimate = –156, SE = 29, t_[df]_ = –5.44_[33]_, p = 5.1 × 10^-6^), with the estimate indicating that lacosamide is associated with a reduction in the SDTC compared to the placebo. The interaction term further reveals that the relationship between SDTC and rheobase differs with lacosamide compared to placebo (estimate = 20.6, SE = 3.9, t_[df]_ = 5.28_[33]_, p = 8.1 × 10^-6^). Specifically, the effect of rheobase on SDTC is less pronounced with lacosamide. For carbamazepine, the results suggest a trend towards an increase in the SDTC (estimate = 84.2, SE = 41.9, t_[df]_ = 2_[33]_, p = 0.053) and a more negative relationship between rheobase and SDTC compared to placebo (estimate = –13.9, SE = 6.8, t_[df]_ = –2.06_[33]_, p = 0.047). However, the effects were considerably smaller in magnitude than those associated with lacosamide.

### 3.5. Evaluation of input-output curve slope

The *k* parameter, representing the slope, displayed the expected strength-duration behaviour^31,32^ in each experimental condition, with higher *k* values for longer pulse durations compared to shorter ones (Figure 5). This was confirmed by the main effect of pulse width in the ANOVA within the LME model framework (F_[df]_ = 34.8_[2, 108]_, p = 2.12 ×10^-12^), with the fixed effects showing that 60 µs (estimate = 0.065, SE = 0.012, t_[df]_ = –5.35_[108]_, p = 4.97×10^−7^, Cohen’s *d* = 2.34) and 120 µs pulses (estimate = 0.132, SE = 0.016, t_[df]_ = 8.34_[108]_, p = 2.65×10^−1^^3^, Cohen’s *d* = 4.76) exhibited greater *k* compared to the reference 30 µs pulses. However, there was no main effect of medication on *k* (F_[df]_ = 0.72_[2, 108]_, p = 0.488) or any interaction of pulse width and medication condition (F_[df]_ = 0.987_[4, 108]_, p = 0.418).

**Figure 5.**
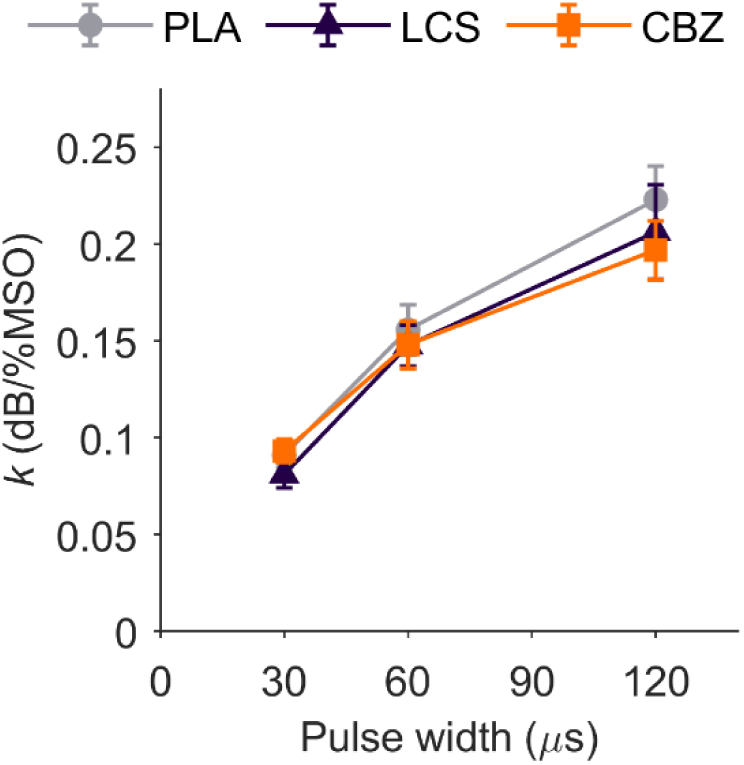
Slope (*k*) of the input-output curve for each condition (placebo, carbamazepine, lacosamide) and pulse width (30, 60, 120 μs), estimated from the log-transformed MEP amplitudes and corresponding stimulus intensities. The *k* parameter reflects the steepness of the curve, with higher values indicating a steeper slope. Analyses showed that *k* was greater for 60 µs and 120 µs pulses compared to 30 µs pulses, but no significant effects of medication condition were observed.

## 4. DISCUSSION

We used a novel TMS device to study the effects of sodium channel-blocking medications on the strength–duration behaviour of neurones in the human M1. Both carbamazepine and lacosamide increased motor thresholds, reducing cortical excitability in line with previous studies using a single pulse width^10–12^. However, single-threshold measurements alone could not distinguish between the effects of the two medications. By employing strength–duration curves, we uncovered distinct pulse-width-dependent effects: carbamazepine increased thresholds proportionally across all pulse widths, while lacosamide’s effects were proportionally larger at longer-duration pulses. These differences were further reflected in the SDTC and rheobase parameters, with lacosamide reducing SDTC, increasing rheobase, and altering their relationship slope, whereas carbamazepine produced only minor and inconsistent effects. Our findings demonstrate the utility of strength–duration metrics in uncovering subtle medication-specific effects on VGSC function, which may be overlooked by single pulse width approaches.

### 4.1. Motor thresholds and I-O curve slope

The observed increases in motor thresholds were not accompanied by changes in I-O curve slopes, aligning with prior studies that reported no effects of sodium channel blockers on MEP amplitudes at fixed intensities relative to motor threshold^10–12^. These findings indicate that the distribution of axonal excitability remains unchanged, and points to uniform effects across axonal populations.

The motor threshold is directly proportional to the magnitude of membrane depolarisation of the neurones stimulated by TMS^32^, which depends on the stimulus strength and duration. However, since TMS activates corticospinal neurones trans-synaptically, changes in a single motor threshold may involve synaptic as well as axonal contributions, making it challenging to isolate the cause. Strength–duration curves, derived from thresholds across multiple pulse widths, address this limitation by focusing on axonal membrane changes. As pulse width increases, the degree of membrane depolarisation also increases, lowering the motor threshold. This shapes the strength–duration curve, which primarily reflects axonal excitability, as axonal depolarisation precedes synaptic activity summation. As such, strength duration curves offer a more precise measure than a single threshold.

### 4.2. TMS-derived strength–duration curves as indices of axonal excitability

Recent work suggests that the cortical SDTC is sensitive to the TMS stimulus waveform, potentially reflecting differential polarisation of the axonal membrane and variations in sodium conductance^30^. This reinforces the idea that strength–duration curves are specific to axonal properties and can serve as indicators of VGSC-mediated changes in cortical axonal excitability. Computational modelling also points to the axon, likely near the terminal, as the site of activation for cortical neurones^7,40^, further supporting the interpretation that strength–duration curves reflect sodium channel changes at these sites.

### 4.3. Medication-induced changes in strength–duration behaviour

Lacosamide significantly altered strength–duration parameters, increasing rheobase and reducing SDTC, highlighting their potential as biomarkers for medication action. These effects mirror those observed in peripheral nerve SDTC^25^. While peripheral changes are plausible, the present findings are more likely to reflect cortical effects. This is because the shape of the strength–duration curve primarily reflects excitability of axons depolarised directly by the TMS pulse, upstream of both cortical synaptic transmission and peripheral motor nerve activation (see Section 4.1). As such, peripheral changes are unlikely to account for the effects observed here.

In contrast, carbamazepine had minimal impact on SDTC or rheobase despite reducing cortical excitability. Notably, preliminary findings from a study indicated that carbamazepine altered the SDTC and rheobase of peripheral sensory nerves in both humans and rodents but not in human motor nerves, despite producing changes in other excitability measures^41^. Our findings regarding motor cortical axons align with the behaviour observed in peripheral motor axons.

The SDTC is influenced by resting sodium conductance, membrane potential, and passive axonal properties like myelination^21–23^. Previous studies of human peripheral nerve excitability after short-term administration with lacosamide^25^ and long-term administration of mexiletine^26^ attributed SDTC increases to changes in sodium conductance. Further evidence comes from cases of accidental tetrodotoxin ingestion, where the potent sodium channel blocker reduced the SDTC^42^. Modelling of the toxin-induced changes in axonal excitability supported VGSC blockade as the likely cause. We therefore interpret our results as being consistent with the proposed VGSC-blocking action of lacosamide.

Membrane potential shifts could also theoretically explain lacosamide-induced changes in strength–duration behaviour^39,43^. However, as noted previously^44^, the low slopes of the relationships between membrane potential and both SDTC and rheobase^43,45^ suggests that a large hyperpolarising shift would be needed to explain changes in strength–duration behaviour.

The differential effects of carbamazepine and lacosamide suggest distinct mechanisms of action, although we can only speculate on the specifics. Axonal membranes contain VGSCs that conduct transient and persistent sodium currents, both of which influence the SDTC, though the latter is more commonly associated with it ^22,23,30,46^. *In vitro* experiments show that carbamazepine and lacosamide reduce both transient and persistent sodium currents^47–50^ [reviewed in^6^]. However, work in mice has shown that carbamazepine induces a hyperpolarising shift in the activation of persistent sodium currents^50^. This produced a significant increase in persistent sodium current conductance at subthreshold voltages in Scn1b knockout mice, which lack the β1 subunit, but not in wild-type mice. This genotype-specific effect could, in some cases depending on β subunit composition, reduce the drug’s effectiveness at suppressing the persistent currents that influence the SDTC. Another relevant factor is that motor axons are known to have lower SDTCs than sensory axons, potentially reflecting reduced persistent sodium currents^23^. These factors may together explain the lack of effect of carbamazepine on SDTC in both peripheral motor axons^41^ and cortical motor axons in the present study.

Differences between the medications may also stem from their interactions with different VGSC gating states. Carbamazepine binds preferentially to VGSCs in the fast inactivated state, delaying recovery from this state^6,47,48^. Meanwhile, lacosamide enhances entry into the slow inactivated state and slows recovery from it^6,47,48^. The latter observation aligns with our findings that lacosamide reduced the SDTC, as the persistent sodium currents involved are prone to slow inactivation^51^, though we note that there is indirect evidence of carbamazepine affecting slowly inactivating sodium currents^52^.

Finally, carbamazepine may exert multiple actions that can interact with or mask its effect on VGSCs, such as its influence on GABAergic synaptic transmission. One TMS study reported effects on cortical silent periods mediated by GABA-B receptors^10^. However, these findings have not been consistently replicated^11^, and the effects on GABAergic inhibition remain unexplored in the present study. Interestingly, it has been suggested that the strength–duration behaviour of excitatory and inhibitory neurones may differ^53^, which could complicate the evaluation of GABAergic inhibition, as TMS intensities are typically set relative to motor threshold. One solution would be to simultaneously examine the strength–duration behaviour of both excitatory and inhibitory synaptic inputs. This approach would benefit the study of novel sodium channel blockers that selectively target VGSC isoforms differentially expressed in excitatory and inhibitory neurones^54^, by providing evidence of effective and selective targeting.

### 4.4. Directions for Future Research

In this study, we aimed to confirm the involvement of axonal membrane-bound VGSCs in the observed medication effects on cortical excitability. Future research could further validate the specificity of SDTC by examining the effects of medications not expected to influence VGSCs, such as those that act primarily at synapses, like levetiracetam^14,20^.

The current TMS-EMG approach is limited to investigating axonal excitability within M1. The development of equivalent TMS-EEG protocols could allow similar measures to be obtained from non-motor regions, potentially enabling the study of region-specific pathophysiology and pharmacological responses, for example, in focal epilepsies. Notably, we have previously shown that TMS-evoked EEG responses are sensitive to stimulation pulse width^55^, suggesting that pulse-width–dependent metrics analogous to SDTC may be technically feasible with TMS-EEG.

Strength–duration metrics may ultimately provide a valuable tool for predicting treatment response, monitoring efficacy, and guiding dose titration in a data-driven, personalised manner. By assessing these metrics in patients initiating therapy, particularly in conditions involving sodium channel dysfunction, clinicians may be able to move beyond trial-and-error approaches toward more targeted and efficient management. Future studies should evaluate the clinical utility of this approach.

## 5. CONCLUSIONS

Our findings highlight distinct effects of sodium channel-blocking medications on strength– duration behaviour in human M1 neurones, with lacosamide aligning with axonal sodium conductance blockade and carbamazepine showing more complex effects. These results underscore the need to consider medication-specific VGSC interactions to improve mechanistic understanding. Strength–duration metrics could advance TMS applications in studying channelopathies, monitoring clinical outcomes, and evaluating the impact of precision-targeted VGSC blockers.

## Funding

R.H.’s contribution was supported by the Academy of Medical Sciences Springboard scheme (SBF009\1073), which is funded by British Heart Foundation, Diabetes UK, the Government Department for Science, Innovation and Technology (DSIT) and Wellcome. A.V.P.’s contribution to the reported research was supported by the National Institutes of Health under Award Numbers RF1MH124943, R01NS117405, and R01MH128422. The content is solely the responsibility of the authors and does not necessarily represent the official views of the funding agencies.

## Acknowledgements

We would like to thank Brainbox Ltd. for the use of the TMS device.

## Declaration of interests

A.V.P. is an inventor on patents on TMS technology and has received: consulting fees and patent royalties for a license on the cTMS (Elevate TMS) technology used in this study from Rogue Research; equity options, scientific advisory board membership, and consulting fees from Ampa Health; consulting fees from Magnetic Tides and Soterix Medical; equipment loan from MagVenture; and research funding from Motif. The other authors have no relevant disclosures.

## Data availability

All data and code used for analysis and figure generation are freely accessible at the following link: https://osf.io/d6vf2/.

## Notes

https://osf.io/d6vf2/

