## Supplemental Data for "Distinct impacts of sodium channel blockers on the strength-duration properties of human motor cortex neurones"

### SUPPLEMENTARY FIGURES

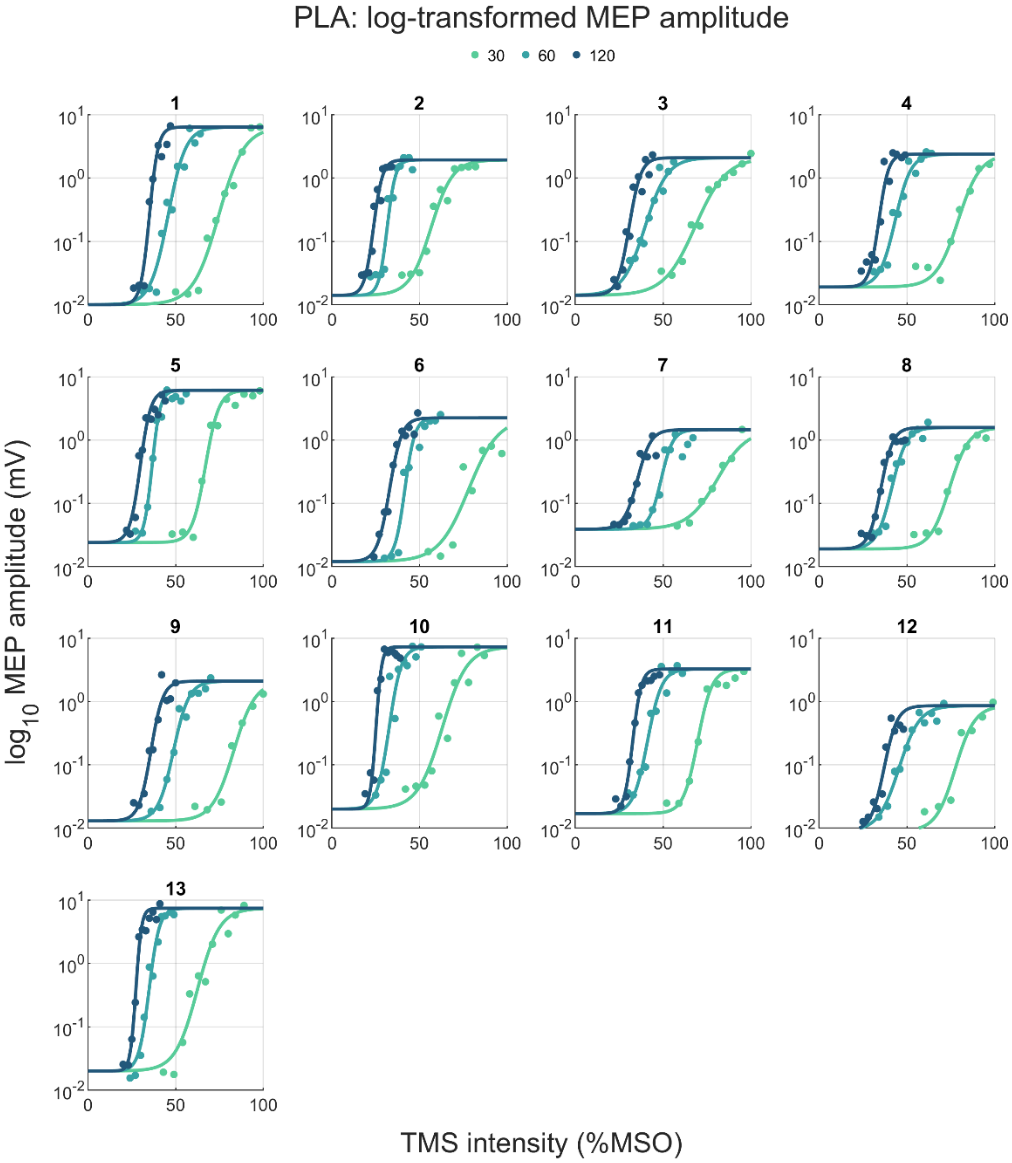


**Figure S1.** I-O curves for each pulse width from each participant in the placebo condition. Data reflect the median log-transformed MEP peak-to-peak amplitude as a function of TMS stimulus intensity.

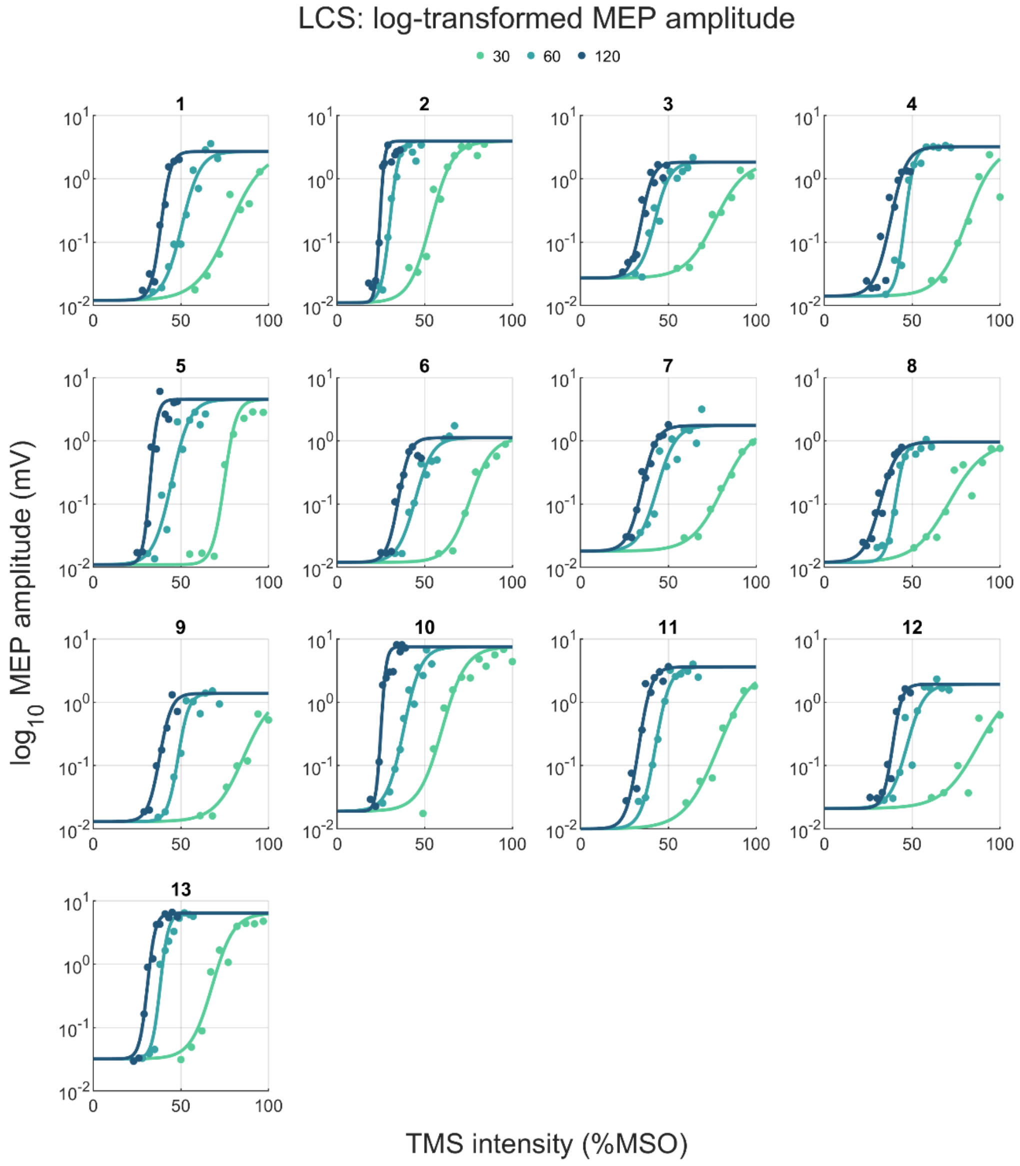


**Figure S2.** I-O curves for each pulse width from each participant in the lacosamide condition. Data reflect the median log-transformed MEP peak-to-peak amplitude as a function of TMS stimulus intensity.

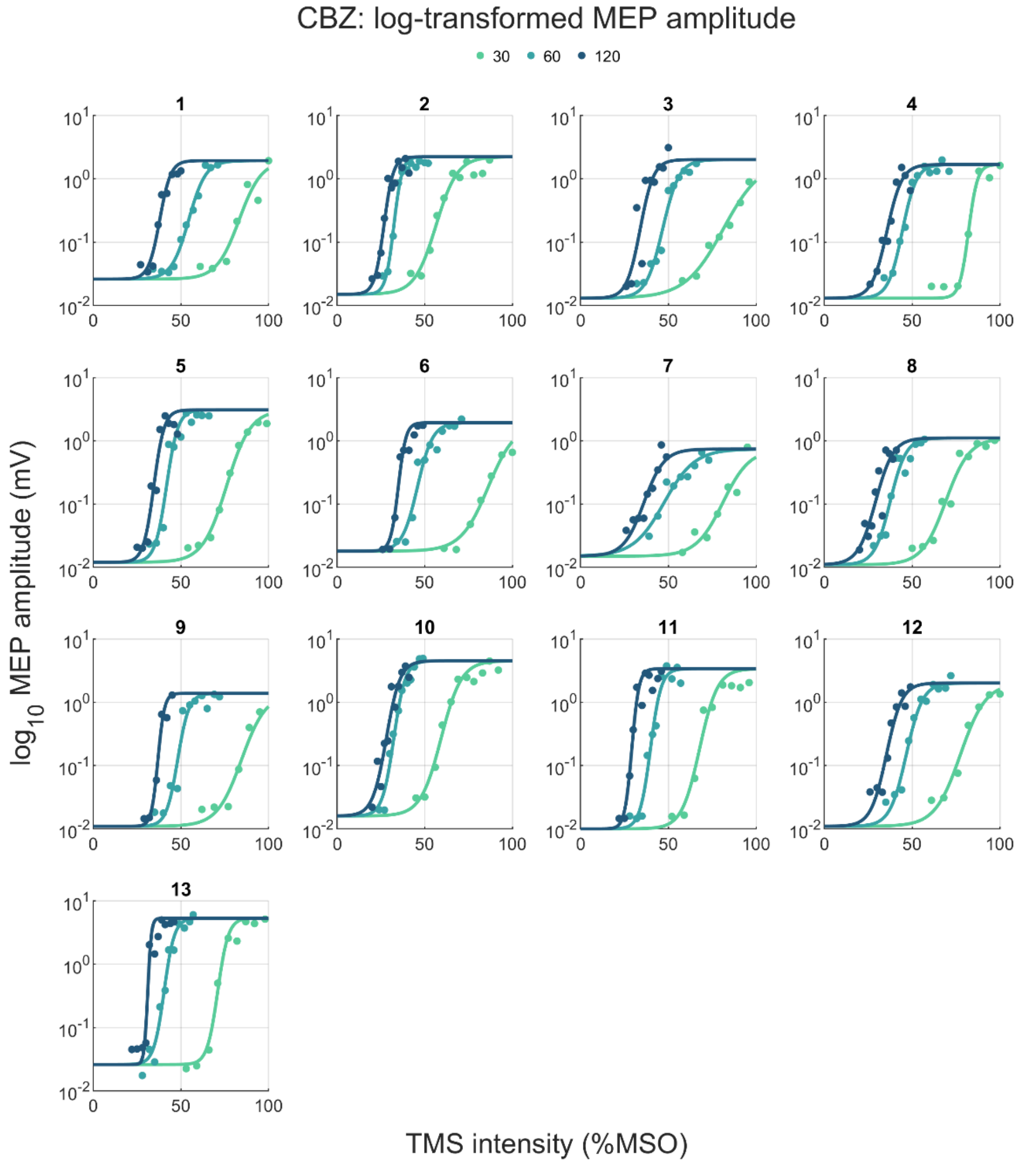


**Figure S3.** I-O curves for each pulse width from each participant in the carbamazepine condition. Data reflect the median log-transformed MEP peak-to-peak amplitude as a function of TMS stimulus intensity.

#### Evaluation of strength-duration time constant under the assumption of a common SDTC

We analysed the data under the assumption of a common SDTC for all participants within the same medication intervention condition but with individual rheobase values. This approach was adopted to avoid potential overfitting issues associated with estimating SDTC separately for each participant[1]. However, since the model assumes a single common SDTC for each medication condition, it is not possible to perform statistical comparisons of SDTC values across different medication conditions directly. To address this limitation, we employed a bootstrapping technique to generate a distribution of SDTC values for comparison purposes.

In this bootstrapping process, we randomly sampled with replacement the paired data points ($RMT_{I\text{-}O}$ and pulse width) from the group within each medication condition. For each bootstrap iteration, we randomly selected pairs of $RMT_{I\text{-}O}$ and pulse width values from the original dataset of participants, allowing for the possibility of repeated selections of the same pair. We then estimated the common SDTC and individual rheobase values from the resampled data. To generate a single value for each iteration, we averaged the rheobase values across all participants. This resulted in one SDTC and one averaged rheobase value per iteration. This procedure was repeated across 1000 bootstrap iterations to create empirical distributions of the SDTC and averaged rheobase values. The estimates from each iteration formed the basis of our distributions of SDTC and rheobase, facilitating comparison across different medication conditions.

To evaluate statistical differences between conditions when assuming a common SDTC across participants, we employed a bootstrap-based t-test approach. We first verified the normality of the data using a Kolmogorov-Smirnov test. For each parameter (SDTC and rheobase), we calculated the bootstrapped paired differences between medication conditions and applied a t-test to these differences to assess statistical significance. To quantify the magnitude of the differences, we calculated Cohen's $d$ for each comparison, defined as the mean of the bootstrapped differences divided by their standard deviation. To estimate the uncertainty around the effect size, we used bootstrapping to derive confidence intervals for Cohen's $d$. We resampled the differences 1000 times, calculating Cohen's $d$ for each bootstrap sample. The 95% confidence interval was obtained by determining the 2.5th and 97.5th percentiles of the bootstrapped effect sizes. This interval provides a range within which the true effect size is likely to fall.

The mean bootstrapped estimates for SDTC and rheobase (Table S1), as well as the overall pattern of differences between conditions under the common SDTC assumption, were broadly comparable to those obtained when assuming individual SDTC values (Figure 4A and 4B). This suggests that the average parameter estimates are robust regardless of the fitting approach.

Statistical analysis on the bootstrapped data confirmed the findings. The bootstrapped distributions conformed to normality, as assessed by a Kolmogorov-Smirnov test, validating the use of parametric tests. We conducted paired t-tests on 1000 bootstrap samples to evaluate the differences between conditions. Significant differences were observed across all comparisons for both SDTC and rheobase (Table S1). However, the differences between placebo and carbamazepine were small in absolute terms and associated with small-moderate effect sizes and are therefore unlikely to be of practical significance. In contrast, comparisons involving lacosamide versus both placebo and carbamazepine showed much larger differences and effect sizes for both SDTC and rheobase.

**Table S1.** Summary of bootstrapped data and statistical analyses (t-test and Cohen's d) for SDTC and rheobase across medication conditions (placebo, PLA; carbamazepine, CBZ; lacosamide, LCS)

| | **Mean ± Bootstrap SEM** | **Mean difference [95% CI]** | $t_{df}$ | **p** | ***d* [95% CI]** |
|---|---|---|---|---|---|
| **SDTC (µs)** | | | | | |
| PLA vs. CBZ | 254 ± 26 vs. 273 ± 22 | -19 [-21, -17] | $-17.3_{999}$ | **$p = 1.4 \times 10^{-58}$** | -0.54 [-0.62, -0.48] |
| PLA vs. LCS | 254 ± 26 vs. 204 ± 16 | 51 [49, 53] | $50.9_{999}$ | **$p = 1.1 \times 10^{-279}$** | 1.61 [1.52, 1.71] |
| CBZ vs. LCS | 273 ± 22 vs. 204 ± 16 | 69 [58, 71] | $80.6_{999}$ | **p = 0#** | 2.55 [2.53, 2.68] |
| **Rheobase (%MSO)** | | | | | |
| PLA vs. CBZ | 6.1 ± 0.6 vs. 6.1 ± 0.4 | -0.10 [-0.05, 0.04] | $-0.2_{999}$ | 0.81 | -0.01 [-0.07,0.05] |
| PLA vs. LCS | 6.1 ± 0.6 vs. 7.8 ± 0.6 | -1.65 [-1.70, -1.59] | $-60.2._{999}$ | **p = 0#** | -1.91 [-2.02, -1.80] |
| CBZ vs. LCS | 6.1 ± 0.6 vs. 7.8 ± 0.6 | -1.64 [-1.69, -1.60] | $-76.3_{999}$ | **p = 0#** | -2.42 [-2.53, -2.31] |
